# Simultaneous dimensionality reduction and integration for single-cell ATAC-seq data using deep learning

**DOI:** 10.1101/2021.05.11.443540

**Authors:** Wolfgang Kopp, Altuna Akalin, Uwe Ohler

## Abstract

Advances in single-cell technologies enable the routine interrogation of chromatin accessibility for tens of thousands of single cells, shedding light on gene regulatory processes at an unprecedented resolution. Meanwhile, size, sparsity and high dimensionality of the resulting data continue to pose challenges for its computational analysis, and specifically the integration of data from different sources. We have developed a dedicated computational approach, a variational auto-encoder using a noise model specifically designed for single-cell ATAC-seq data, which facilitates simultaneous dimensionality reduction and batch correction via an adversarial learning strategy. We showcase both its individual advantages on carefully chosen real and simulated data sets, as well as the benefits for detailed cell type characterization via integrating multiple complex datasets.

## Main

Rapid advancements in single-cell epigenomics technologies, including single-cell ATAC-seq (scATAC-seq), have enabled the interrogation of gene regulation at an unprecedented resolution. scATAC profiles the accessibility of chromatin across the whole genome and is the currently most widely adapted protocol to identify candidates of regulatory regions of importance to the system under investigation. The resulting datasets require specialized computational tools to cope with their characteristic high dimensionality and sparsity and will rely on scalability for future datasets.

A key step in every scATAC-seq processing pipeline is dimensionality reduction, which aims to represent the most salient trends in the data at lower dimensionality, such as groups of similar cells. The quality of this step is critical, as it precedes other analysis tasks, including cell-type characterization, identifying cell-type specific regulatory regions, motif analysis, etc. Several methods have been introduced for dimensionality reduction using scATAC-seq data, including latent Dirichlet allocation (cisTopic) [1], latent Semantic indexing (LSI) [2], SnapATAC [3] and SCALE [4]. A recent benchmark analysis showed that these tools work well for cell type characterization for small or moderate dataset sizes, but may not scale to large dataset sizes and/or vary in performance across different datasets [5]. Apart from performing dimensionality reduction, the growing number of published datasets opens up new avenues for data integration of replicates or data obtained with different protocols, such as combinatorial indexing or droplet-based approaches [6].

To address the lack of dedicated single-cell ATAC-seq tool that enable simultaneous data integration (e.g., batch correction) and dimensionality reduction, we have developed BAVARIA, a batch-adversarial variational auto-encoder (VAE) [7] that facilitates dimensionality reduction and integration for scATAC-seq data. To this end, we extended the standard VAE framework in several ways: First, inspired by combining deep learning with specialized and suitable count noise models for processing single-cell RNA-seq data (e.g., by using a zero-inflated negative binomial distribution [8, 9]), we set out to find a suitable count model for scATAC-seq. We identified a reconstruction loss that is based on the negative multinomial distribution, which allows to model the raw accessibility profile as a whole instead of considering regions independently and account for over-dispersed distributions of the count data. This negative multinomial loss outperforms alternative reconstruction loss functions with respect to the quality of the latent features for cell-type characterization, including the binary-cross-entropy or the multinomial-derived loss (see Suppl. Fig. 1; Methods). Second, fitting neural networks is commonly based on stochastic optimization, which may lead to variable latent feature quality across multiple training runs. Due to the optimizer getting stuck in a poor local optimum, cell-types may occasionally be poorly characterized. Here, an ensemble approach, whereby latent features of several independently trained models are concatenated, not only stabilizes the latent feature quality, but also appears to improve their feature quality compared to latent features from individual models (see Suppl. Fig. 2; Methods). Third, we adopted a domain-adversarial training strategy [10] that encourages the VAE to extract latent features uninformative of batch effects. Specifically, we use batch-discriminator networks not only at the final layer of the encoder as suggested in Ganin et al. [10], but also at intermediate layers of the encoder (see Fig. 1A). This puts more emphasis on removing irrelevant batch-associated information in the initial layers of the network, which we find to enhance the batch correction capabilities (see comparison below and Methods). We refer to our approach as batch-adversarial VAE or BAVARIA (Fig. 1A).

**Figure 1.**
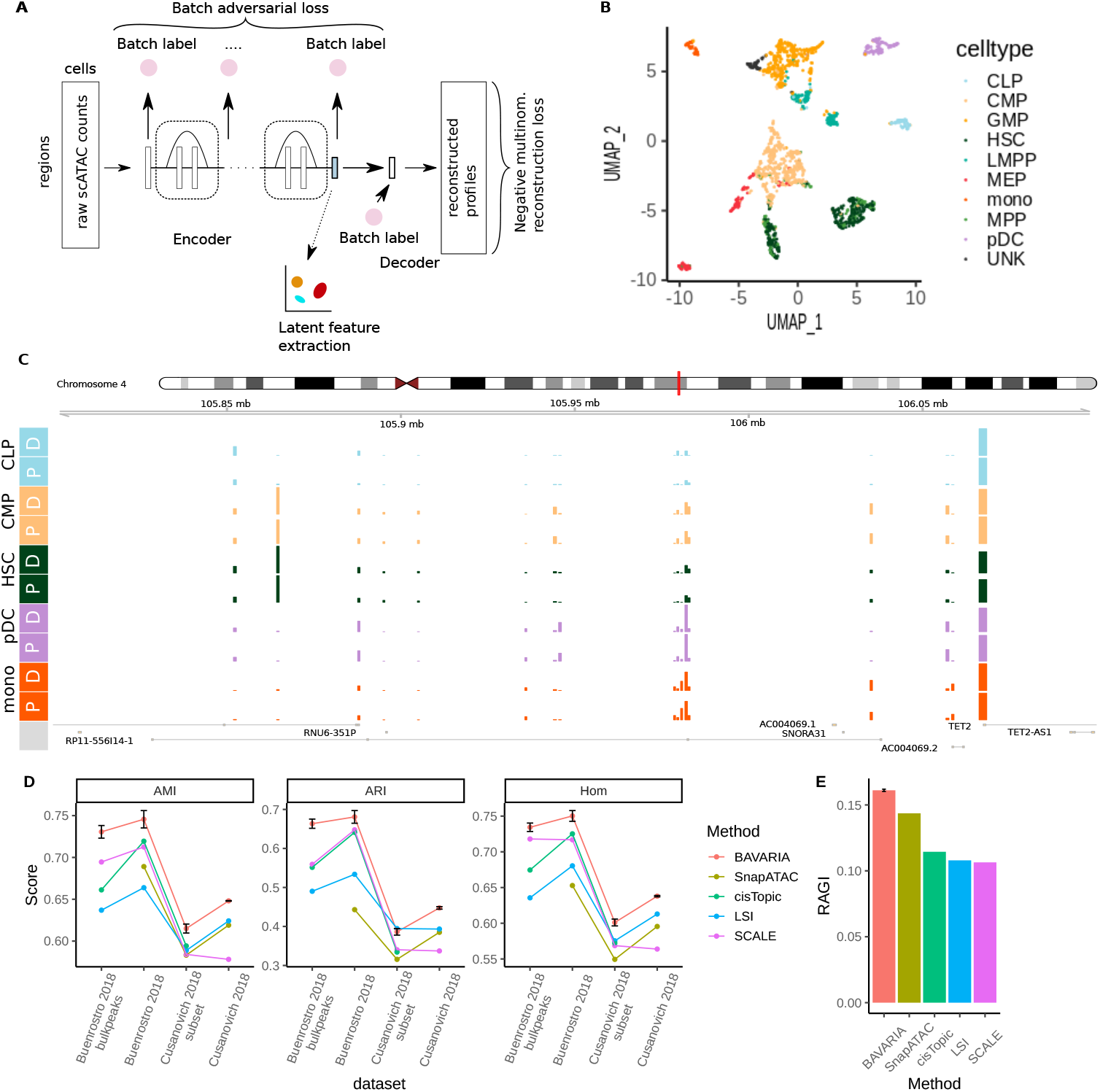
A) Schematic representation of the BAVARIA model, variational auto-encoder implementing a negative multinomial reconstruction loss and batch-adversarial training for batch correction (Batch-Adversarial VARIational Auto-encoder - BAVARIA). Latent features of the encoder network module serve as low-dimensional representation of the high-dimensional original accessibility profile. B) Scatterplot of single cells from hematopoiesis data [12] in the UMAP space. Latent features were obtained by fitting BAVARIA ten times from scratch to mitigate fluctuations due to random initialization and concatenating the latent features of the individual models (excluding outlier models; see Methods). The concatenated latent features were used as input to the UMAP algorithm. C) Comparison between observed and predicted pseudo-bulk accessibility profiles for hematopoiesis data [12]. BAVARIA was fitted ten times from scratch. Predicted single-cell accessibility tracks were averaged across the models (excluding outlier models) and subsequently collapsed within the known cell types CLP, CMP, HSC, pDC and mono cells. Pseudo-bulk tracks for the original input data and the prediction are indicated using the suffix D and P, respectively. D) Cell clustering based on the derived low-dimensional feature representations were evaluated by comparing clusters against known ground truth cell labels on hematopoiesis data (Buenrostro 2018 and Buenrostro 2018 bulkpeaks [12]) and mouse tissue cells (Cusanovich 2018 and Cusanovich 2018 subset [2]) using the adjusted mutual information (AMI), adjusted Rand index (ARI) and homogenity (Hom). Generally, high scores indicate good behavior for capturing known cell populations. Buenrostro 2018 and Buenrostro 2018 bulkpeaks use the same cells but different peak regions [5, 12] and Cusanovich 2018 subset consists of a subset of cells from Cusanovich 2018 [5] (see Methods). Results for LSI, cisTopic and SnapATAC were obtained from the benchmark assessment [5]. An ensemble of BAVARIA was run *n* = 3 times. The performances are summarized by the mean score (dot) and error bars that denote the +/- SEM. E) Clustering performance evaluated on 10X Genomics 5k PBMCs [17] using the residual average Gini index (RAGI) which is based on marker gene separation [5]. Results for LSI, cisTopic and SnapATAC were obtained from the benchmark assessment [5]. An ensemble of BAVARIA was run *n* = 3 times. The performances are summarized by the mean score (dot) and error bars that denote the +/- SEM.

We systematically assessed the ability of BAVARIA derived low-dimensional feature representations for cell type characterization on a range of real and synthetic datasets. To this end, we expanded on a recently published bench-marking framework [5] for comparing to current state-of-the-art solutions (cisTopic [1], LSI [2], SnapATAC [3] and SCALE [4]). Briefly, for each method, a low-dimensional feature representation is extracted and subjected to cell clustering. For BAVARIA, the low-dimensional representation is derived from the latent features of the encoder. Subsequently, a range of clustering evaluation scores are used to determine how well the cell clusters reflect cell identities based on known ground truth cell labels (if available) or by assessing the separation of known marker genes (see Methods and [5]).

Across three publicly available datasets, we observe variable performance of LSI, cisTopic, SnapATAC and SCALE. That is, no single method consistently outperformed the other methods across the datasets (see Fig. 1D,E). By contrast, BAVARIA robustly achieves best or equal performance for uncovering cell identities across all real datasets (see 1D,E) which is also exemplified by the separation of known cell types in the UMAP embedding (see 1B) and the high similarity of network-imputed and original signal tracks within distinct known cell types (see 1C).

Next, we assessed the performance of the methods at different read depths and noise levels using two synthetic datasets (a bonemarrow and an erythropoiesis dataset). We generated larger simulated datasets relative to [5], as we found previous sizes to be insufficiently small to reflect current realistic experiments and for fitting large neural networks (see Methods). For the synthetic bonemarrow datasets, LSI and BAVARIA both achieve similar high performance across all sparsity levels, while the performance of cisTopic, SnapATAC and SCALE decrease sooner with decreasing read depth (see 250 and 500 fragments per cell; see Suppl. Fig. 3A). All methods perform similarly well on higher noise levels for the bonemarrow dataset (see Suppl. Fig. 3B). On a synthetic erythropoiesis dataset, all methods achieve similar results for the high read coverage regime (see 5000 fragments per cell; Suppl. Fig. 3C). Yet, with decreasing read coverage, BAVARIA outperforms the other methods (see 1000, 500 and 250 fragments per cell; Suppl. Fig. 3C). In addition, LSI and BAVARIA perform best for 0% and 20% additional noise, and BAVARIA outperforms all other methods for 40% additional noise using synthetic erythropoiesis data (see Suppl. Fig. 3D).

Having established the superior performance on benchmark tasks, we turned to BAVARIA’s signature feature of data integration. We analyzed two adult mouse brain cell samples from different sources: 10X Genomics [11] and Cusanovich et al. 2018 [2] (see Methods). To quantify the contribution of our new data integration strategy, we first disabled adversarial training with BAVARIA. Here, cells from the two datasets occupy non-overlapping territories in the cell embedding space, which underlines the severity of the batch effect (see Fig. 2A). That is, cells largely cluster by batches. By contrast, with batch-adversarial training, BAVARIA achieves a markedly better alignment between cells from different batches, while also largely maintaining a separation between previously annotated cell types (see Fig. 2B).

**Figure 2.**
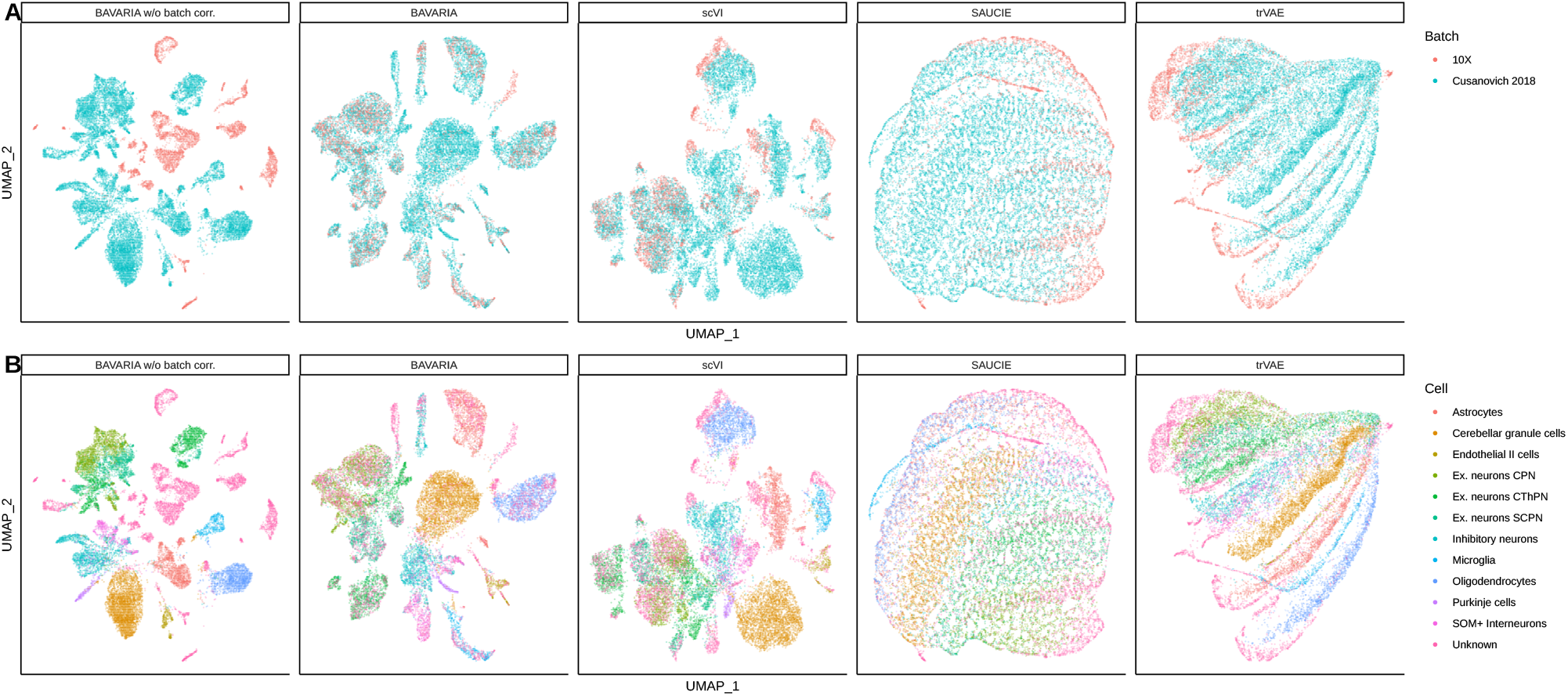
A) UMAP embedding illustrating cells from 10X Genomics and Cusanovich et al. 2018 after applying BAVARIA w/o batch correction, BAVARIA, scVI [9], SAUCIE [19] and trVAE [18]. B) UMAP embedding illustrating previously characterized cell types [2] (Astrocytes, Cerebellar granule cells, Encothelial II cells, Ex. neurons CPN, Ex. neurons CThPN, Ex. neurons SCPN, Inhibitory neurons, Microglia, Oligodendrocytes, Purkinje cells, SOM+ interneurons and unknown cells). 10X Genomics cells [11] are labelled ‘Unknown’ as no labels are avaible.

Since current dimensionality reduction tools do not offer the ability of simultaneous data integration, we decided to compared BAVARIA against three other deep learning models that enable batch integration (scVI, trVAE and SAUCIE) but were originally developed for scRNA-seq data. This enabled us to assess whether a dedicated approach to the characteristics of single-cell open chromatin would indeed surpass a naive strategy to use a tool tailored to a different modality. Compared to scVI, trVAE and SAUCIE, As expected, BAVARIA indeed achieves a better separation between cell clusters (see Fig. 2B), since the other tools were developed for processing scRNA-seq data. However, it underscores the importance of developing specialized tools for different single-cell data modalities. In addition, residual batch effects can still be appreciated after batch correction for scVI, SAUCIE and trVAE (see Fig. 2A). Likewise, we also observe residual batch effects when using a conditional VAE variant of BAVARIA (similar as proposed for scVI [9]) and when applying batch-adversarial training using only a single batch-discriminator network at the final layer of the encoder (as in Ganin et al.; see Suppl. Fig. 4).

Next, we clustered the cells based on the joint latent features and inspected pseudo-bulk accessibility profiles stratified by the batches around several marker genes. For clusters with relatively even representation of cells from both batches, highly concordant accessibility profiles across clusters can be observed (e.g., Scl17a7, Caln1, Gad2, Tmem119, Aldh1l1, Mbp and Abca4; Suppl. Fig. 5A,B,D,E). This suggests that the latent features derived from BAVARIA are suitable for cell label transfer, as cells appear to cluster together based on their underlying cell type. On the other hand, as a sign for successful integration of data taken from similar but not identical sources, we also find cell clusters that are primarily present in one of the samples. For instance, cluster 0 and 18 consist of mostly Cusanovich et al. 2018 cells which exhibit accessibility at Mmp24 and Itga11, whereas cluster 16 consists of mostly 10X Genomics cells which exhibit cluster-specific accessibility around Sh3bp4. While label transfer from an annotated reference onto a dataset without cell-type annotation is possible in a supervised manner (e.g., by classifying cell types based on their accessibility profile), unsupervised data integration (e.g., via BAVARIA) offers the possibility to correct and account for imperfections of the original cell annotations (e.g., reference dataset). For instance, we observe several sub-populations of cells that have previously been annotated as inhibitory neurons (e.g., cluster 4 and 14; see Suppl. Fig. 5A). Similarly, a cell population previously annotated as unknown (cluster 22; Suppl. Fig. 5C) likely represents doublet events between cerebellar granular cells and oligodendrocytes, as these cells exhibits specific accessibility for both cell types (compare cluster 0, 2 and 22; Suppl. Fig. 5C).

In summary, we have developed a VAE for integrating sparse and high-dimensional scATAC-seq data (BAVARIA). We demonstrated that several unique aspects, regarding robust training and noise modelling for scATAC count data, allow the model to match and exceed the performance of current state-of-the-art solutions for cell type characterization across 1) different dataset sizes, 2) different read depths and 3) different noise levels. Importantly, its batch-adversarial training strategy makes BAVARIA the first tool to facilitate data integration and accurate batch correction across different scATAC protocols.

## Methods

### Data preparation - benchmark analysis

Data for the benchmark analysis was obtained by following and adapting a recently published scATAC-seq benchmarking framework [5]. We obtained 1) a hematopoietic differentiation dataset (n=2034 cells; Buenrostro 2018; [12]), 2) a single-nucleus combinatorial indexing ATAC-seq mouse tissue dataset (n=81173 cells; Cusanovich 2018; [2]) and 3) a peripheral blood mononuclear cell dataset (n=5335 cells; 10x PBMC 5k). The former two datasets also include ground truth cell labels from FACS sorted cells or by using the tissue of origin as label. The benchmark also includes a 15% subset of cells from the mouse tissue dataset (n=12178 cells; Cusanovich 2018 subset; [2]).

The hematopoietic dataset was independently processed twice with two different sets of peaks. Namely, peaks determined on the single-cell ATAC-seq by Chen et al. 2019 [5] (Buenrostro 2018) and the original peak set reported by Buenrostro et al. 2018 [12] (Buenrostro 2018 bulkpeaks).

Preprocessing includes feature counting in the pre-defined peak regions, binarization of the count matrix, filtering for a minimum read coverage per peak (at least 1% of cells need to be covered at the region), and removing sex chromosomes. This led to 98738, 132110, 378894, 141388 and 67427 peaks for the Buenrostro 2018, Buenrostro 2018 bulkpeaks, Cusanovich 2018, Cusanovich 2018 subset and 10x datasets, respectively.

We followed the procedure of Chen et al. 2019 [5] to generate several synthetic datasets based on FACS-sorted bulk-ATAC-seq samples from bone-marrow [13] and erythropoiesis [14], as described previously [5].. As the originally published benchmarking assessment consists of too few cells for fitting large neural networks, we increased the numbers of cells. Specifically, for bonemarrow, 2000 cells per population were generated with different fragment sizes per cell (5000, 2500, 1000, 500 and 250 fragments) as well as for different noise levels (0%, 20% and 40% additional noisy reads). Likewise, for erythropoiesis, 1000 cells per population were generated with different fragment sizes per cell (5000, 2500, 1000, 500 and 250 fragments) as well as for different noise levels (0%, 20% and 40% noise). Note also that the downsampling experiment based on the erythropoiesis data was not part of the original benchmarking assessment [5]. Finally, for all synthetic datasets, the 80000 most covered regions were retained for the benchmark analysis.

### Data preparation - mouse brain cell integration

We downloaded fresh adult mouse brain cell data from the 10X Genomics web site (https://support.10xgenomics.com/single-cell-atac/datasets/1.2.0/atac_v1_adult_brain_fresh_5k) as well as single-cell ATAC-seq data from several mouse tissues [2] from https://atlas.gs.washington.edu/mouse-atac/data/.

We used the peaks provided by 10X Genomics as reference peaks [11]. The master peaks from Cusanovich et al. 2018 were lifted over from mm9 to mm10 and mapped onto the reference peaks using bedtools intersect [15]. Likewise, the original count matrix was mapped onto the new reference peak set and only cells from the WholeBrain and PreFrontalCortex tissues were retained for the analysis [2]. The count matrices from the 10X Genomics and Cusanovich et al. 2018 datasets were concatenated, binarized and filtered to ensure that each region was covered in at least 1% of the cells. Regions on the sex chromosomes were removed. The final count matrix contained 18605 cells (3880 from 10x and 14725 from Cusanovich et al.) and a peak set of size 136528.

### Negative multinomial variational auto-encoder

We define the accessibility profile of a given single cell across a set of regions as *x* = [*x*_1_, *…, x*_*N*_] where *x*_*i*_ > 0 reflects accessibility of region i. *x*_*i*_ = 0 indicates inaccessibility. In the context of this work, we use binarized accessibility profiles (e.g., *x*_*i*_ = 0 or *x*_*i*_ = 1), although the model is in general also applicable to non-negative integer vectors (e.g., sites with multiple reads per cell).

In this section, we first introduce the adaptation of the standard VAE model [7] without integrated batch-correction and turn to batch-adversarial training strategy in the next section. Briefly, following Kingma and Welling 2014 [7], the encoder determines the *L*-dimensional mean *µ* and variance *σ* parameters of the approximate Gaussian posterior distribution of the latent feature representation given an accessibility profile *x*. From the approximate posterior distribution, samples *z* are drawn, which are in turn used as input for the decoder to reconstruct the *N* dimensional accessibility profile [7]. We shall use the mean vector *µ* as the low-dimensional feature representation used for the downstream analysis.

We assume that the accessibility profile *x* follows a negative multinomial distribution defined by

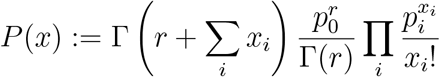

where *p* = [*p*_0_, …, *p*_*N*_] represent the non-negative parameters which sum to one and *r* denotes the positive real-valued dispersion parameter. *p*_*i*_ for *i* > 0 reflects the accessibility profile, while *p*_0_ is associated with the dispersion.

We construct a decoder that determines the respective parameters *p* and *r*. In particular, for *i* > 0 the decoder computes

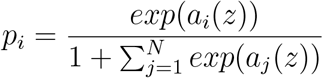

where *a*_*i*_ represents the decoder’s output activity at region *i* for a given latent feature sample *z*. The remaining probability mass is reserved for the dispersion and is given by

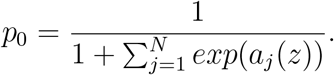

Here we assume that the dispersion parameter is scalar and does not depend on the accessibility profile *x*, i.e., it is adjusted like a bias term in the network.

Consequently, the reconstruction loss is given by

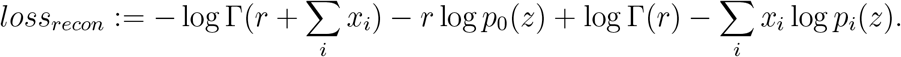

As described previously [7], Kullback-Leibler divergence is utilized as a regularization loss, denoted as *loss*_*KL*_, which encourages that the latent feature representation is distributed according to 𝒩(0, *I*). The total loss is given by summing the reconstruction loss and the regularization loss, which is subject to minimization by adapting the model parameters during the model fitting.

### Batch-adversarial training

To facilitate data integration and batch correction, we took inspiration from Ganin et al. 2016 [10] and adapted the variational auto-encoder framework described above to enable batch-adversarial training. The goal of this approach is to establish a latent feature representation that captures biologically relevant information about the accessibility profiles (e.g., to describe cell types), while at the same time conveying as little information as possible about experimental batch labels.

To facilitate batch-adversarial training, the standard VAE architecture is augmented by batch-discriminator network modules. These sub-networks are stacked on top of the final layer of the encoder for the purpose of predicting the batch label from the latent feature representation. We use batch-discriminator networks with softmax output activation to predict categorical batch labels. Multiple independent batch labels can be used simultaneously with separate softmax output units for each batch. In addition to the batch-discriminator network at the final layer of the encoder (similar to Ganin et al. 2016 [10]), we also add batch-discriminators on top of the hidden layers of the encoder (after the first hidden layer and after each residual network block). The batch-discriminator network modules at the initial and intermediate layer allow to put more emphasis on removing batch related information early on in the network, and is intended to simplify disentangling batch-related information from the biologically relevant signal throughout the entire encoder network.

Apart from the additional batch-discriminator networks, we modified the decoder to take as input the latent feature encoding *z* as well as the one-hot encoded batch labels. This enables the decoder to recombine the batch information with the (ideally) batch-corrected latent feature representation to compute the reconstruction of the accessibility profiles. This is important, as batch-related information is still used in this way to compute the reconstruction loss.

Finally, we adjust the training objective of the standard VAE as follows: We measure how well batch-labels can be predicted from the latent features of the encoder (e.g., derived from the hidden or final layer) using the categorical cross-entropy loss,

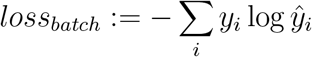

where *y* and *ŷ* denote the true and predicted batch label. While the parameters of the batch-discriminator associated parameters are adapted to minimize *loss*_*batch*_, the parameters associated with the encoder module are adapted according to

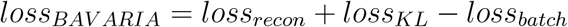

That is, the encoder seeks to find a latent feature representation that is uninformative for the batch-label classification.

### Model and training hyper-parameters

We use the following model architecture for all experiments: For the encoder, we use a feed forward layer with 512 nodes and ReLU activation, followed by 20 consecutive residual neural network blocks with 512 nodes, which feed into two layers representing the means and variances of the latent features (dimensions listed in Suppl. Table 1). Each residual block is composed of a feed forward layer with rectified linear units (ReLU) and a feed forward layer whose output is added to the block’s input before applying ReLU activation. For the decoder, we use a single feed forward layer consisting of 16 and 25 neurons in the benchmarking analysis and the data integration use case, respectively.

For the batch-adversarial training, batch discriminator network modules are stacked on top of the intermediate layers of the encoder (after the first layer and after each residual block) as well as on top of the final layer of the encoder. Each discriminator network consists of two layers with 128 neurons and ReLU activation and an output layer with softmax activation.

The models were fitted using 85% of the cells using AMSgrad [16]. The remaining 15% cells were used for validation. Additional dataset-specific hyper-parameters are listed in Suppl. Table 1.

### Ensemble of models and feature extraction

We fitted a BAVARIA model *M* times, each time starting from random initial weights. Afterwards, we concatenated the mean vectors of the approximate posterior distribution of the latent features *µ* either across all *M* individual models or by using a subset of these models, dependent on the use case. In the latter case, we sought to remove potentially poor quality models whose average loss across the dataset exceeded an outlier criterion. Specifically, we removed models if their average loss after training exceeded *Q*_75%_(*loss*)+1.5*× IQR*(*loss*), where *Q*_75%_ denotes the 75% quantile of the loss distribution across the *M* individual models and IQR represents its interquartile range. The latent features of the remaining models were concatenated and considered for the downstream cell clustering analysis.

### Benchmark analysis

We adapted a recently published scATAC-seq benchmarking framework [5]. For all real datasets, we obtained the results of cisTopic [1], LSI [2] and SnapATAC [3] as previously reported [5]. In addition, we ran LSI on the full Cusanovich 2018 dataset in the same way it was previously run for the Cusanovich 2018 subset [5]. We fitted BAVARIA models on the filtered count matrices using the hyper-parameters described in Suppl. Table 1. After training each individual model, an ensemble model was created by concatenating individual models by excluding outlier models (see above). Latent features of the ensemble were used for the downstream clustering using the benchmarking framework. We ran SCALE [4] with default parameters using the input count matrices described above (e.g., the same matrices which were used for BAVARIA) and extracted the latent features for the down-stream clustering analysis.

For the synthetic datasets, we ran cisTopic, LSI and SnapATAC with the same parameters as in the published benchmark analysis, but using larger numbers of cells. We applied SCALE with parameters ‘–min_cells 0 –min_peaks 0.0’ to ensure that the simulated count matrix is not further subjected to filtering. BAVARIA ensembles were fitted for each synthetic dataset using hyper-parameters listed in Suppl. Table 1.

Following Chen et al. 2019 [5], the latent features extracted for each method were subjected to k-means clustering, hierarchical clustering (with ward linkage), and Louvain clustering. For all datasets with available ground truth labels, the adjusted Rand index (ARI), adjusted mutual information (AMI) and homogeneity (Hom) were used to score the agreement of the clustering results with the known cell labels [5]. For the 10X dataset (without ground truth) [17], the residual average Gini index (RAGI) was computed based on a previously reported set of marker and house keeping genes [5].

Subsequently, the maximum clustering score across the clustering algorithms (k-means clustering, hierarchical clustering or Louvain clustering) was reported for each dataset and score (ARI, AMI, Hom).

### Comparison of alternative reconstruction losses

We compared several reconstruction loss implementations on Buenrostro et al. 2018 data using the benchmarking framework described above [5]. Specifically, we used the same variational auto-encoder architecture parameters, with exception of the final layer of the decoder. Assuming the accessibility profiles are derived from a Bernoulli distribution, we used a sigmoid output activation for the decoder in conjunction with a binary cross-entropy loss, *loss*_*Bin*_ = −Σ_*i*_ (*x*_*i*_ log(*p*_*i*_(*x*)) + (1−*x*_*i*_) log(1− *p*_*i*_(*x*))). Alternatively, assuming multinomially distributed accessibility profiles, we used a softmax output activation in conjunction with a negative log-likelihood of the multinomial distribution, *loss*_*Mul*_ = −Σ _*i*_ *x*_*i*_ log *p*_*i*_(*x*).

### Data integration of mouse brain cells

We used BAVARIA with a 15 dimensional latent feature representation and 25 neurons for the hidden layer of the decoder. We trained an ensemble of 10 individual models for 200 epochs using a batch size of 64 (Suppl. Table 1). In addition, each individual model was fitted on a random 50% subset of the original peak set, which enabled training time speedup while maintaining similar clustering qualities (data not shown).

Due to the lack of dimensionality reduction tools that support data integration for single-cell ATAC-seq, we compared BAVARIA to several tools that were originally established for single-cell RNA-seq analysis and facilitate simultaneous dimensionality reduction and batch correction: We fitted scVI [9] with default parameters. We trained trVAE [18] by following the tutorial code (https://nbviewer.jupyter.org/github/theislab/trVAE/blob/master/examples/trVAE_Haber.ipynb) using the unnormalized count matrix (136528 regions by 18605 cells). We fitted SAUCIE [19] with default parameters.

As baseline, we ran simplified versions of the BAVARIA architecture: 1) we adjusted BAVARIA to a conditional VAE model, which receives the batch labels as input at the encoder (along with the raw read profile), rather than predicting the batch labels from the latent features and hidden layers. 2) We adjusted BAVARIA to use only a single batch-discriminator network module at the final layer of the encoder (along the lines of Ganin et al. 2016; [10]). These networks were trained with the same network parameters and hyper-parameters as BAVARIA.

The UMAP embedding was computed based on the latent features from each method using scanpy [20]. For the UMAP visualization, cells previously annotated as Astrocytes, Cerebellar granule cells, Encothelial II cells, Ex. neurons CPN, Ex. neurons CThPN, Ex. neurons SCPN, Inhibitory neurons, Microglia, Oligodendrocytes, Purkinje cells, SOM+ interneurons and unknown cells in Cusanvovich et al. 2018 and all 10X Genomics cells [11] were illustrated.

### Clustering and differential accessibility analysis of mouse brain data

Using the BAVARIA derived latent features, we performed Louvain clustering using scanpy [20]. Cluster-specific accessibility was determined by using a generalized linear model and a binomial distribution:

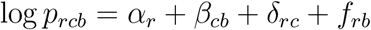

where *α*_*r*_ denotes the region-specific offset for region *r, β*_*cb*_ denotes the cluster and batch-specific offset for cluster *c* and batch *b, δ*_*rc*_ denotes the cluster-specific accessibility for region *r* and cluster *c* and *f*_*rb*_ denotes the batch-specific accessibility for region *r* and batch *b*.

After fitting the linear models, we identified the 100 top most accessible regions per cluster by ranking *δ*_*rc*_ and visualized the associated raw accessibility profiles in a heatmap using scanpy [20].

For the pseudo-bulk visualization, we re-mapped reads from Cusanovich et al. 2018 WholeBrain and PreFrontalCortex tissues to mm10 using bowtie2 [21] using the parameters ‘–very-sensitive -X 2000 -3 1’. Cluster-specific pseudobulk bam files were constructed by dividing the reads by barcodes associated with the clusters. These bam files were converted to bigwig tracks using ‘bamCoverage -normalizeUsing CPM’ from deeptools [22]. Finally, pyGenomeTracks [23] was used to visualize the cluster-specific accessibility tracks.

## Data availability

All datasets to perform the benchmark analysis were obtain from the computational scATAC-benchmarking framework https://github.com/pinellolab/scATAC-benchmarking. This includes the publicly available single-cell ATAC-seq 10X PBMC 5k dataset [17], the hematopoiesis dataset [12] and the adult mouse dataset [2], as well as the bonemarrow [13] and erythropoiesis dataset [14] from which the simulated single-cell ATAC-seq datasets were derived. For the brain data integration, we additionally obtained mouse brain cells from Cusanovich et al. [2] https://atlas.gs.washington.edu/mouse-atac/data/ and the 10X Genomics website https://support.10xgenomics.com/single-cell-atac/datasets/1.2.0/atac_v1_adult_brain_fresh_5k.

## Code availability

BAVARIA is available via GitHub under a GPL-v3 license at https://github.com/BIMSBbioinfo/bavaria

## Acknowledgements

The authors wish to thank Pia Rautenstrauch, Jonathan Ronen and Remo Monti for valuable comments on the manuscript. This work was supported by the German Federal Ministry of Education and Research (de.NBI; FKZ 031L0101B) and by the Helmholtz Association (sparse2big; ZT-I-0007).

## Author Contributions

