## Supplementary Information for "Simultaneous dimensionality reduction and integration for single-cell ATAC-seq data using deep learning"

Kopp *et al.*

Supplementary Table 1: Summary of hyperparameters for the experiments. Number of replications refers to the number of re-trained models with random initial weights. $*$ denotes the dataset and setup on which the binary and multinomial noise models were evaluated.

| Dataset | Epochs | Batch size | latent dims / hidden layer dims. | Ensemble size |
|---|---|---|---|---|
| Buenrostro 2018* | 100 | 64 | 10/16 | 10 |
| Buenrostro 2018 bulkpeak | 100 | 64 | 10/16 | 10 |
| Cusanovich 2018 subset | 120 | 256 | 10/16 | 10 |
| Cusanovich 2018 full | 100 | 256 | 30/16 | 3 |
| 10x 5k PBMC | 100 | 64 | 10/16 | 10 |
| Bonemarrow clean | 100 | 128 | 3/16 | 10 |
| Bonemarrow coverage 5000 | 100 | 128 | 3/16 | 10 |
| Bonemarrow coverage 2500 | 100 | 128 | 3/16 | 10 |
| Bonemarrow coverage 1000 | 100 | 256 | 3/16 | 10 |
| Bonemarrow coverage 500 | 200 | 512 | 3/16 | 10 |
| Bonemarrow coverage 250 | 150 | 512 | 3/16 | 10 |
| Bonemarrow coverage 20% noise | 100 | 256 | 3/16 | 10 |
| Bonemarrow coverage 40% noise | 200 | 512 | 3/16 | 10 |
| Eryhropoiesis clean | 100 | 256 | 3/16 | 10 |
| Eryhropoiesis 20% noise | 100 | 256 | 3/16 | 10 |
| Eryhropoiesis 40% noise | 100 | 256 | 3/16 | 10 |
| mouse brain cell integration | 200 | 64 | 15/25 | 10 |

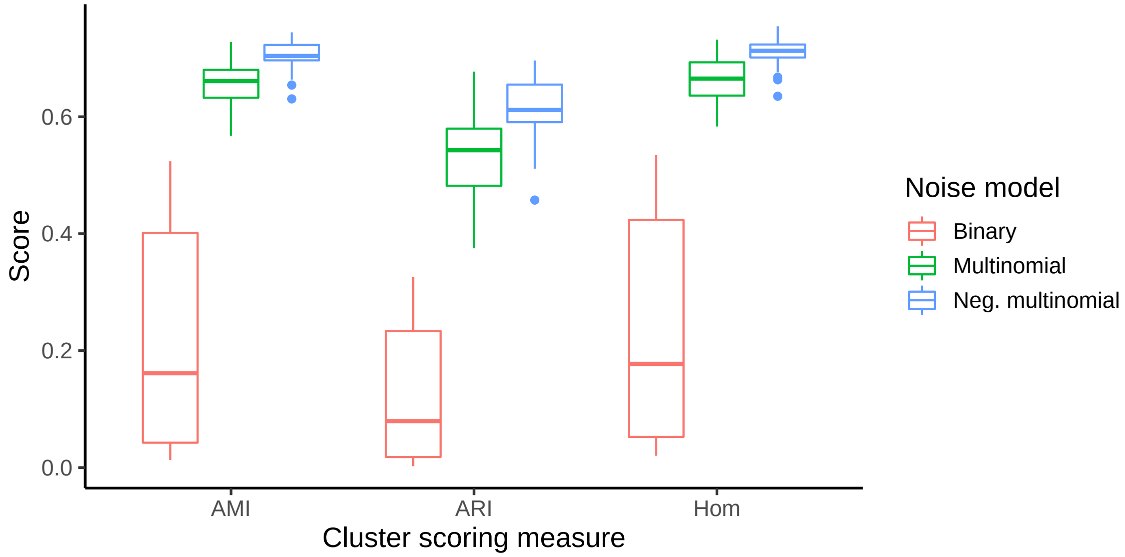


Supplementary Figure 1: **Comparison of reconstruction loss measures.** The suitability of different reconstruction loss measures was assessed by fitting thirty individual models on the Buenrostro *et al* 2018 dataset. The total loss across the dataset was determined for each model and models with poor outlier losses were excluded (e.g. due to poor local minima; see Methods), leading to 28, 29 and 29 models for binary, multinomial and negative multinomial loss for the visualization, respectively. The x-axis represents different reconstruction losses: binary cross-entropy loss (Binary), negative log-likelihood for the multinomial distribution (Multinomial), and negative log-likelihood of the negative multinomial (Neg. multinomial) distribution. Otherwise, the model architecture remained the same. Latent features were subjected to clustering using k-means, hierarchical clustering and Louvain clustering and clustering performances were computed based on adjusted mutual information (AMI), adjusted Rand index (ARI) and Homogeneity (Hom) against ground truth cell labels. The best score across the clustering algorithms was considered. Boxes represent quartiles Q1 (25% quantile), Q2 (median) and Q3 (75% quantile); whiskers comprise data points that are within 1.5 x IQR (inter-quartile region) of the boxes.

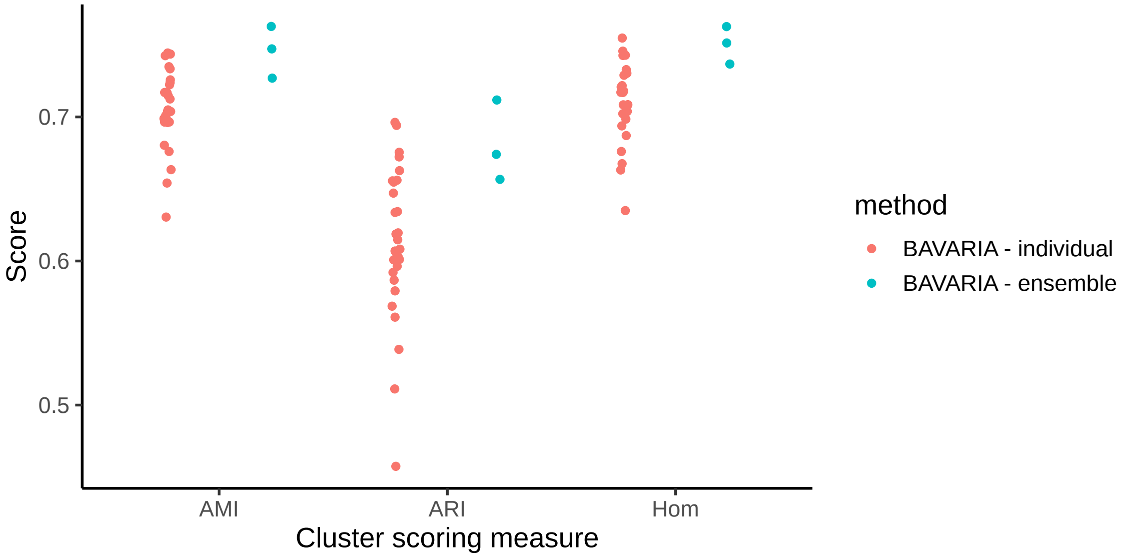


Supplementary Figure 2: **Combining latent features of separately trained models.** Three ensembles consisting of ten VAE models were fitted on the Buenrostro et al. 2018 dataset. The total loss across the dataset was determined for each model and models with poor outlier losses were excluded from the ensemble (e.g., due to poor local minima; see Methods). Individual models (BAVARIA - individual) were combined to ensembles by concatenating the latent features (BAVARIA - ensemble). Latent features were subjected to clustering using several algorithms and clustering performances were computed based on adjusted mutual information (AMI), adjusted Rand index (ARI) and Homogeneity (Hom). The best score across the clustering algorithms are considered.

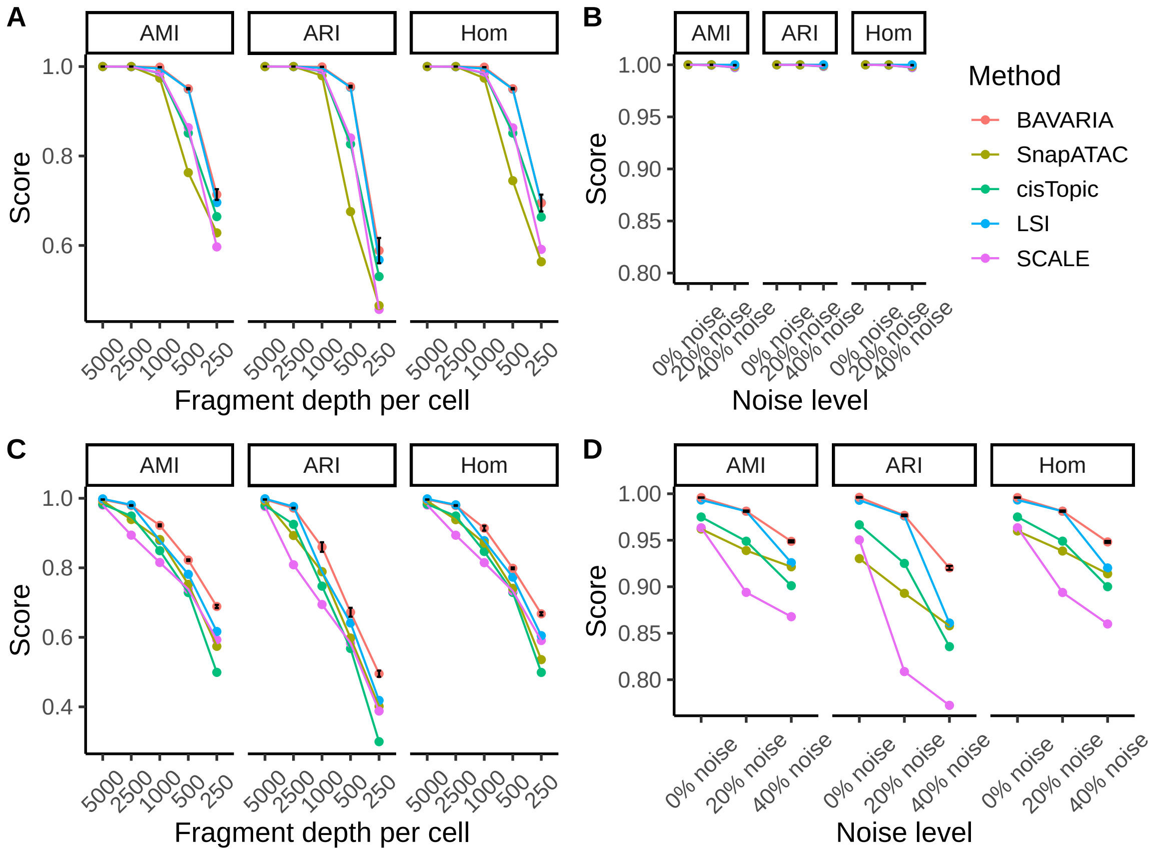


Supplementary Figure 3: **Cell type characterization assessment using synthetic data.** A) Bonemarrow data using 5000, 2500 1000, 500 and 250 fragments per cell. B) Bonemarrow data using 0%, 20% and 40% additional noise. C) Erythropoiesis data using 5000, 2500 1000, 500 and 250 fragments per cell. D) Erythropoiesis data using 0%, 20% and 40% additional noise. Low-dimensional feature representations were obtained using cisTopic, LSI, SnapATAC, SCALE and BAVARIA and subjected to clustering using different algorithms (k-means, hierarchical clustering, Louvain clustering). Clustering performances were evaluated using adjusted mutual information (AMI), adjusted Rand index (ARI) and Homogeneity (Hom) compared against ground truth cell labels (see Methods). The best score across clustering algorithms is shown. cisTopic, LSI, SnapATAC and SCALE were run once per case, while $N = 3$ ensembles of NM-VAE were trained from scratch to assess the variability of the performance. The dot represents the mean performance and the error bars indicate the +/- SEM according to the repetitions.

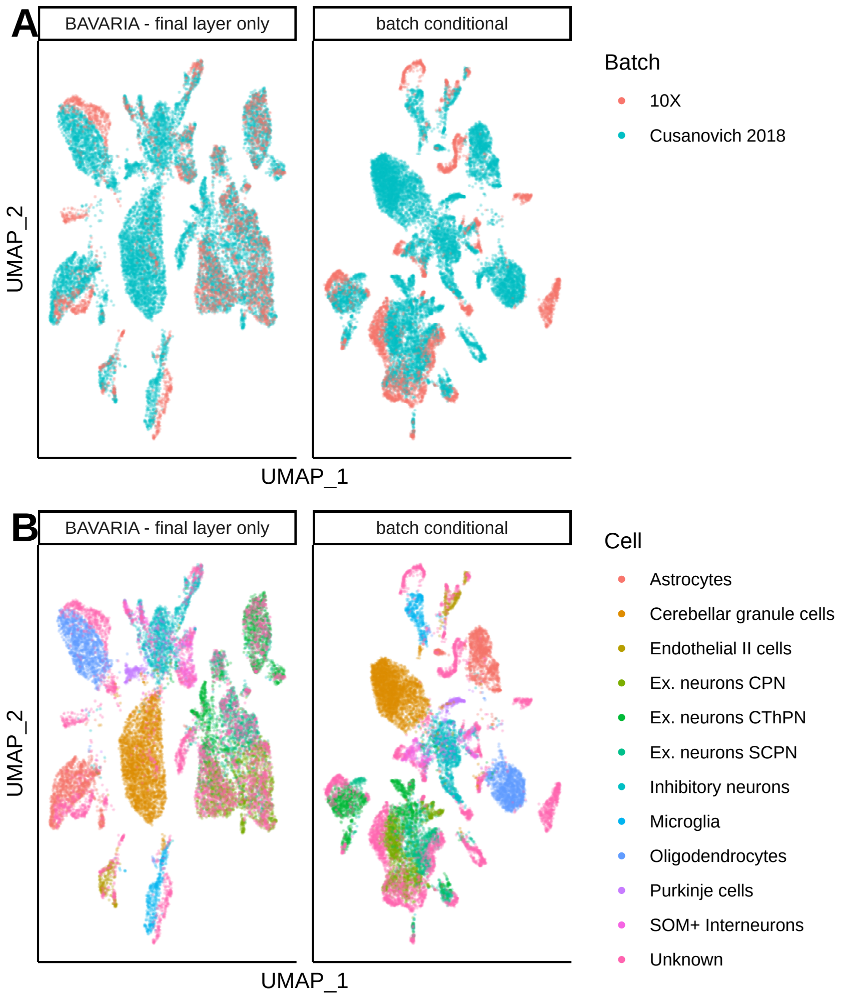


Supplementary Figure 4: **Batch correction - Comparison of architectures.** A) UMAP embedding illustrating cells from 10X Genomics and Cusanovich et al. 2018 after applying a BAVARIA variant with a single batch-discriminator network module at the final encoder layer (BAVARIA - final layer only) and a conditional variational auto-encoder variant of BAVARIA which receives the batch labels as input for the encoder's initial layer (batch conditional). B) UMAP embedding illustrating previously characterized cell types [1] (Astrocytes, Cerebellar granule cells, Encothelial II cells, Ex. neurons CPN, Ex. neurons CThPN, Ex. neurons SCPN, Inhibitory neurons, Microglia, Oligodendrocytes, Purkinje cells, SOM+ interneurons and unknown cells). 10X Genomics cells are labelled 'Unknown'.

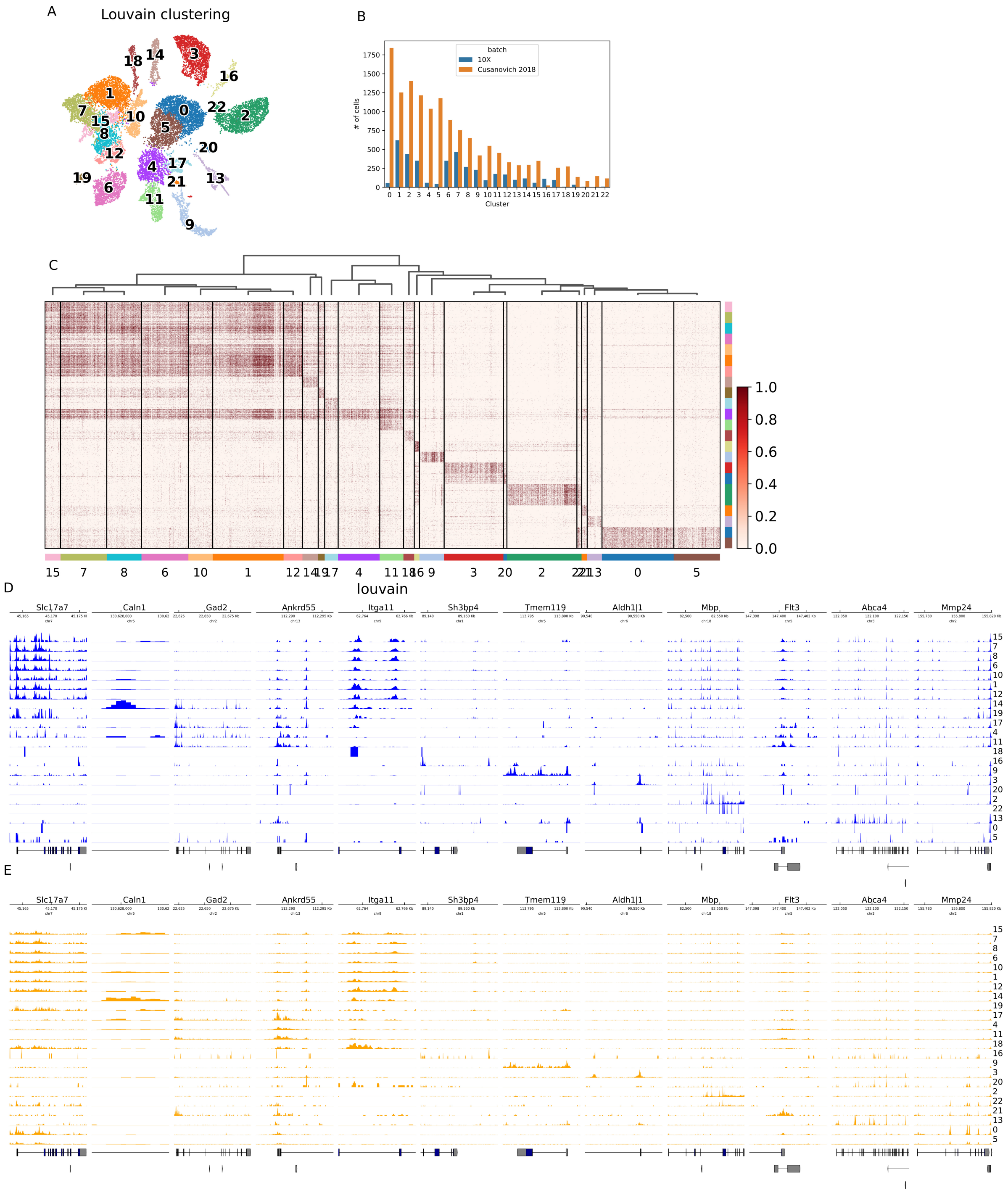


Supplementary Figure 5: **Clustering and cluster-associated regions.** A) Clustering of the integrated 10x and Cusanovich et al. 2018 datasets. B) Number of cells per cluster and batch. C) Illustration of cluster-associated accessibility using the 100 top accessible regions per cluster. D) Depth normalized accessibility tracks per cluster for the 10X dataset for several marker regions. E) Depth normalized accessibility tracks per cluster for the Cusanovich et al. 2018 dataset for several marker regions.
